# Occurrence and Distribution of Common Bacterial Blight and Bacterial Leaf Blight in Tanzanian Lowland and Highland Agro-ecologies using Digital Image Analysis

**DOI:** 10.64898/2025.12.10.693561

**Authors:** Emanuel L. Mulungu, Richard R. Madege, Beatrice Mwaipopo, Michael P. J. Mahenge, Camilius A. Sanga, Farian S. Ishengoma

## Abstract

Common Bacterial Blight (CBB) and Bacterial Leaf Blight (BLB) are important constraints affecting paddy and bean yields, respectively, causing significant losses if unmanaged. For effective planning and management of such diseases, accurate and reliable quantification, i.e., prevalence, incidence, and severity, becomes crucial. Despite this, the quantification of these diseases in Tanzania remains limited. Here, digital image analysis (ImageJ and Plantix) was used in the identification and quantification of CBB and BLB in low and highland agroecologies represented by Kilosa and Mbarali districts, respectively, by surveying 24 paddy and common bean fields across 10 villages. The study revealed that CBB and BLB were highly prevalent (100%) across study sites, with varying incidence and severity. Incidence and severity of both CBB and BLB were significantly higher in Kilosa than in Mbarali, with a large proportion (>19%) of these variations accounted for by district-level differences. Interestingly, significant variations in incidence and severity were observed even at the village level. In Kilosa, CBB severity differed significantly among villages (p = 0.018), while in Mbarali, BLB incidence and severity varied significantly (p < 0.001; p = 0.0436). Village-level differences accounted for over 41% of the total variation. Conclusively, the study indicates that CBB and BLB are highly prevalent across both Kilosa and Mbarali districts and their respective villages, with incidence and severity varying due to both district- and village-level differences. To manage these diseases effectively, site-specific management and geographically targeted interventions are required that account for local variability while prioritizing areas of higher infection risk, such as Kilosa. This approach will not only improve plant disease management programs but also ensure efficient resource use, thereby promoting sustainable disease control under changing climatic conditions.

## Introduction

Paddy and common beans are among the major food crops cultivated in Tanzania, playing a crucial role in sustaining millions of people [1,2]. Despite more than 47% of the country’s land area (44 million hectares) being classified as arable [3], the productivity of these key crops remains well below their potential. In fact, Tanzania currently achieves less than half of the potential yields for both paddy and common beans [4,5]. This sub-optimal productivity may be attributed to various factors, including plant diseases. Among these, blight diseases, namely Common Bacterial Blight (CBB) and Bacterial Leaf Blight (BLB), caused by *Xanthomonas axonopodis* pv. *phaseoli* (*Xap*) and *Xanthomonas oryzae* pv. *oryzae* (*Xoo*), respectively, affect common beans and paddy. CBB has been reported to cause yield losses of up to 40% in susceptible common bean varieties in East Africa, with total crop failure possible if unmanaged [6]. BLB, on the other hand, has been reported in Tanzania to cause paddy yield losses ranging from 13% to 20%, potentially exceeding 60% under favorable conditions [7,8]

Plant disease assessment, also known as phytopathometry, is a fundamental component of effective plant disease management. It involves the identification of plant diseases, followed by the measurement or estimation of the amount of disease expressed as signs or symptoms on single or groups of host plants, using key parameters such as incidence, severity, and prevalence [9,10]. Incidence is defined as the proportion (or percentage) of diseased plants in a whole population of cultivated plants, which is calculated by taking the number of infected plants divided by the total number of plants in the selected population [11]. On the other hand, disease severity is the degree to which a plant or plant part is diseased, given as the area or volume of the plant tissues exhibiting visible disease symptoms [9,11,12], while prevalence is the proportion (or percentage) of number of fields where the disease has been detected at least in one plant or parts of plant [9]. Phytopathometry serves the purpose of not only understanding and monitoring epidemiology, but also further extends its importance to analysis of yield losses caused by diseases, breeding for disease resistance, evaluating and comparing the effectiveness of control methods, all of which are vital for decision making on any plant disease management system [9,13,14].

Phytopathometry, like other pathology disciplines, has advanced significantly due to technological innovations to improve precision and reliability in identifying and quantifying plant diseases. Traditionally, plant disease identification required visual observation, normally by skilled personnel, which is time-consuming and expensive, while disease quantification relied on the raters’ visual observation, often subject to errors and bias [10,13,15]. Among the advancements is the Digital Image Analysis (DIA), which allows the detection, identification, and quantification of diseases in plants or plant parts by extracting the information after analysis of digital images using image analytical tools [9]. Analytical tools such as Plantix and ImageJ are among the most widely used software for plant disease identification and quantification, respectively. These tools have significantly improved the speed and accuracy of disease diagnosis and disease quantification. Plantix can deliver diagnostic results in just a few seconds with an accuracy of 90% to 100%, while ImageJ is simple, easy to use, and enables disease quantification with a relative error as low as 2.9% [16–18].

Despite the aforementioned efficiency of DIA, there is limited use, particularly in providing quantitative measurements on the prevalence, incidence, and severity of plant diseases in Tanzania. As a result, reliable quantification data for CBB and BLB remain inadequate. Thus, the study aimed to establish the status of Common Bacterial Blight (CBB) on common beans and Bacterial Leaf Blight (BLB) on rice. Disease incidence, severity, and prevalence were assessed in two agroecologically distinct areas, the highland agroecological zone represented by Mbarali district in Mbeya region, and the lowland agroecological zone represented by Kilosa district in Morogoro Region, using image analysis software, including Plantix and ImageJ, to obtain accurate diagnoses and quantitative estimates. The study further examined the spatial variability in recorded incidences and severity among the study areas. The results from this study provided baseline information for decision-making on prioritizing plant disease management and for guiding future research aimed at improving current management practices.

The remainder of this paper is structured as follows: Section two presents the materials and methods. Section three outlines the results. Section four provides a discussion of the results. Finally, section five concludes the study with recommendations and limitations.

## Materials and Methods

### Study area

The study was conducted in two distinct agroecological zones, namely, the eastern lowland agroecological zone, represented by Kilosa district (6.8343° S, 36.9917° E), and the southern highland agroecological zone, represented by Mbarali district (8.6713° S, 34.3143° E), as shown in Fig 1., Kilosa, as the representative of lowland agroecological zones, has an elevation ranging from 300 to 600 meters above sea level (a.s.l) [19]. It is one of the nine districts in the Morogoro Region where rice and common beans are the primary crops. In contrast, Mbarali, representing the Highland agroecological zones, has an altitude ranging from 1000 to 1800 meters above sea level [20]. It is one of the seven districts in the Mbeya Region, situated in a lowland semi-arid environment where rice and common beans are the primary crops. The weather conditions, including monthly maximum temperature, minimum temperature, and rainfall in Kilosa and Mbarali districts during the survey period (April and May), are shown in Fig 2.

**Fig 1.**
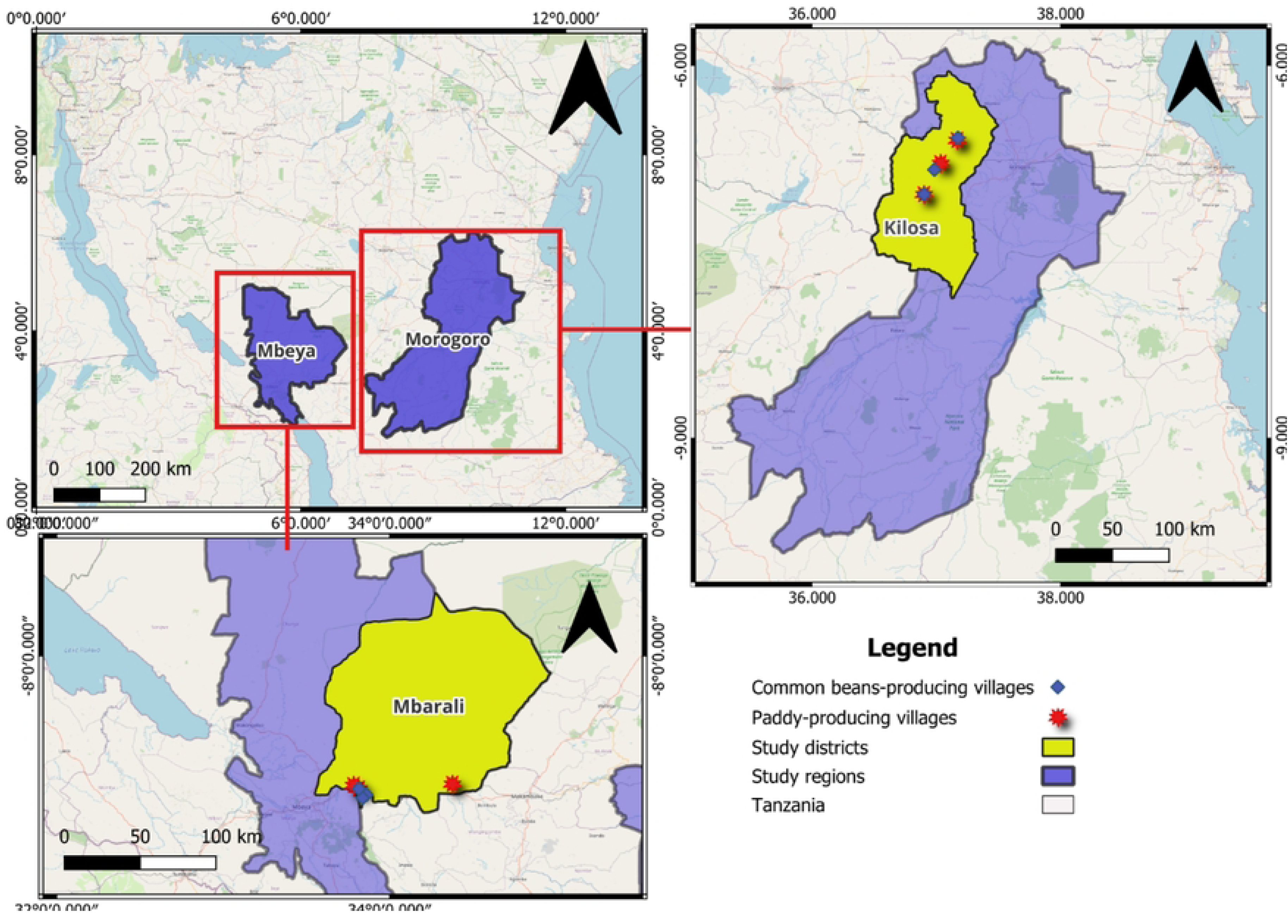
Map showing the locations of the study.

**Fig 2.**
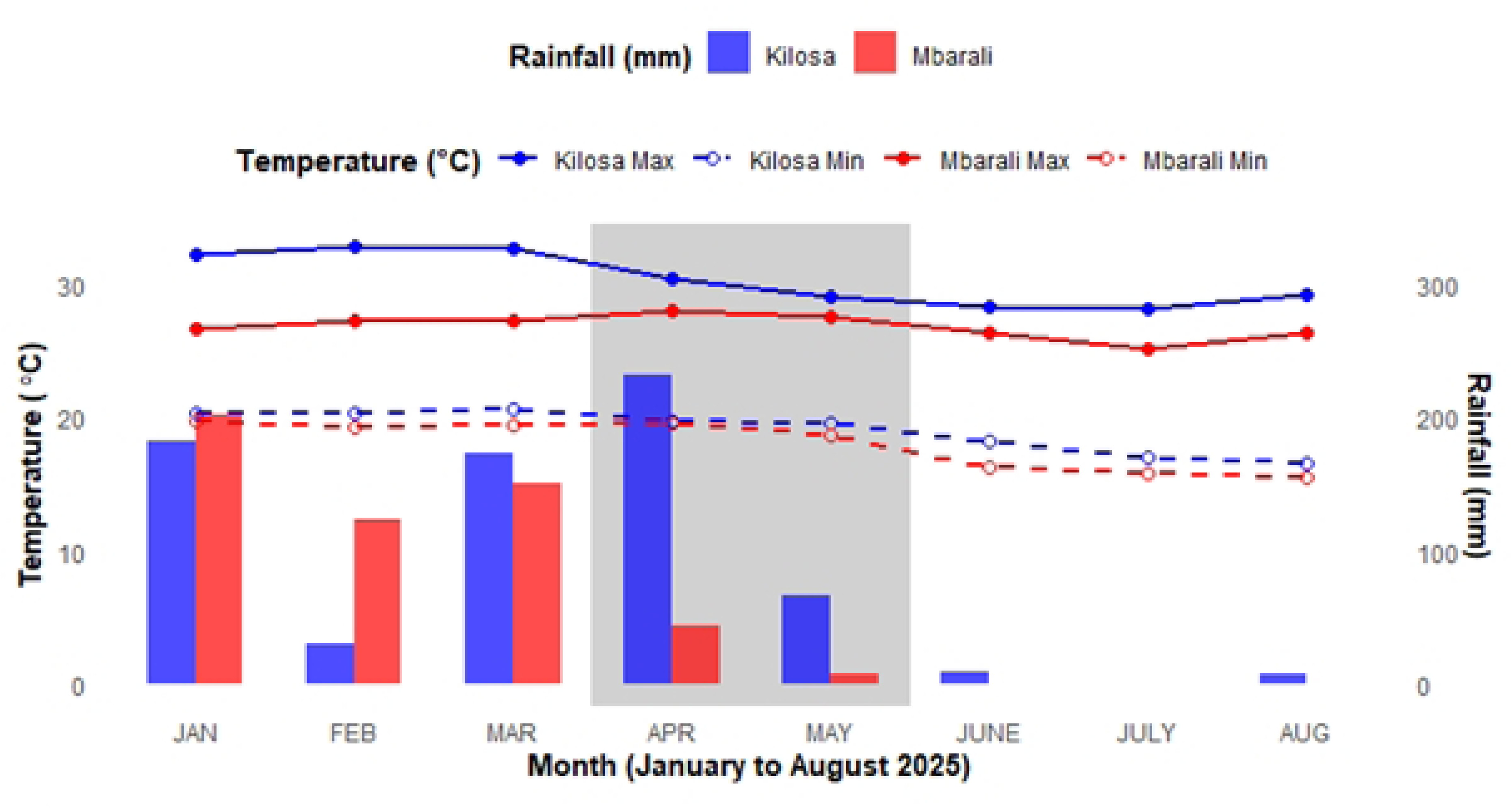
Monthly rainfall (mm) and maximum/minimum temperature (°C) for Kilosa and Mbarali districts, covering January to August 2025. The shaded area indicates the monthly rainfall and temperature (maximum and minimum) conditions during the survey period. Source: Tanzania Meteorological Authority (TMA).

The rationale for choosing the Mbarali and Kilosa districts for the study is that, firstly, Mbarali and Kilosa are among the major producers of rice and common beans, thereby increasing the likelihood of the target diseases being found in these areas. Secondly, weather variability, such as temperature and rainfall, has been reported in these districts, which has also compromised agricultural productivity [21,22]. Thirdly, Kilosa and Mbarali are found within different agroecological zones, with Kilosa found in the lowland agroecology, while Mbarali is found in the highland agroecology; thus, they represent distinct climate profiles that are suitable for examining the influence of spatial variability on plant disease dynamics.

### Study design and Sampling

The study utilized a cross-sectional observational design to quantify CBB and BLB in Kilosa district and Mbarali district at a specific point in time. A purposive multistage sampling method was used to select wards and villages based on the productivity levels of paddy and common beans. With the help of the District Agricultural, Livestock, and Fishery Officer, DALFO (pers. comm.), a total of nine villages from six wards across two districts were selected for the study. In Kilosa, four wards were selected: Chanzulu, Masanze, Mvumi, and Ulaya, comprising five villages: Ilonga-Msalabani, Chabima, Makwambe, Mvumi Gongwe, and Ulaya-Mbuyuni. In Mbarali, two wards were selected: Kongolo-Mswiswi and Igurusi, encompassing five villages: Lunwa, Majenje, Mambi, Kongolo-Mkola, and Azimio-Mswiswi. The field visit was conducted between May 1^st^ and 23^rd^, 2025. A purposive systematic sampling approach was employed to select the surveyed households’ farms based on the following inclusion criteria: a farm with a minimum size of one hectare, cultivating both common beans and paddy, with paddy at the seedling to milking stages and common beans from the seedling stage (V1) to the early pod-filling stage (R8), and located in proximity to motorable roads. The surveys were conducted at regular intervals of 5 to 10 km. A total of 24 household farms were surveyed during the field visit, comprising 12 farms from each study district, with two farms selected per village. Within each household farm, five locations were assessed.

### Assessment of disease incidence, severity, and prevalence

The assessment of disease incidence, severity, and prevalence followed a two-step pipeline, consisting of identification and quantification.

#### a) Identification

Before assessing disease incidence and severity in the field, preliminary identification was conducted on symptomatic plant parts. The identification of CBB and BLB was done using the digital diagnostic tool Plantix (version 4.9.4). Following the method of [23], quadrats were placed in an X-shaped pattern within each field, giving a total of 5 quadrats in each field, and a destructive purposive sampling was used to collect samples of diseased leaves. Only leaves showing symptoms of either CBB in common beans or BLB in paddy were collected, while multi-infected leaves were excluded to avoid errors during identification and subsequent analyses. High-quality images of symptomatic leaves were captured using a Canon PowerShot SX700 HS digital camera (16.1 MP) and analyzed in Plantix, as illustrated in Fig 3. Only plants confirmed positive for CBB or BLB through the software-based diagnosis were included in the subsequent incidence and severity assessments

**Fig 3.**
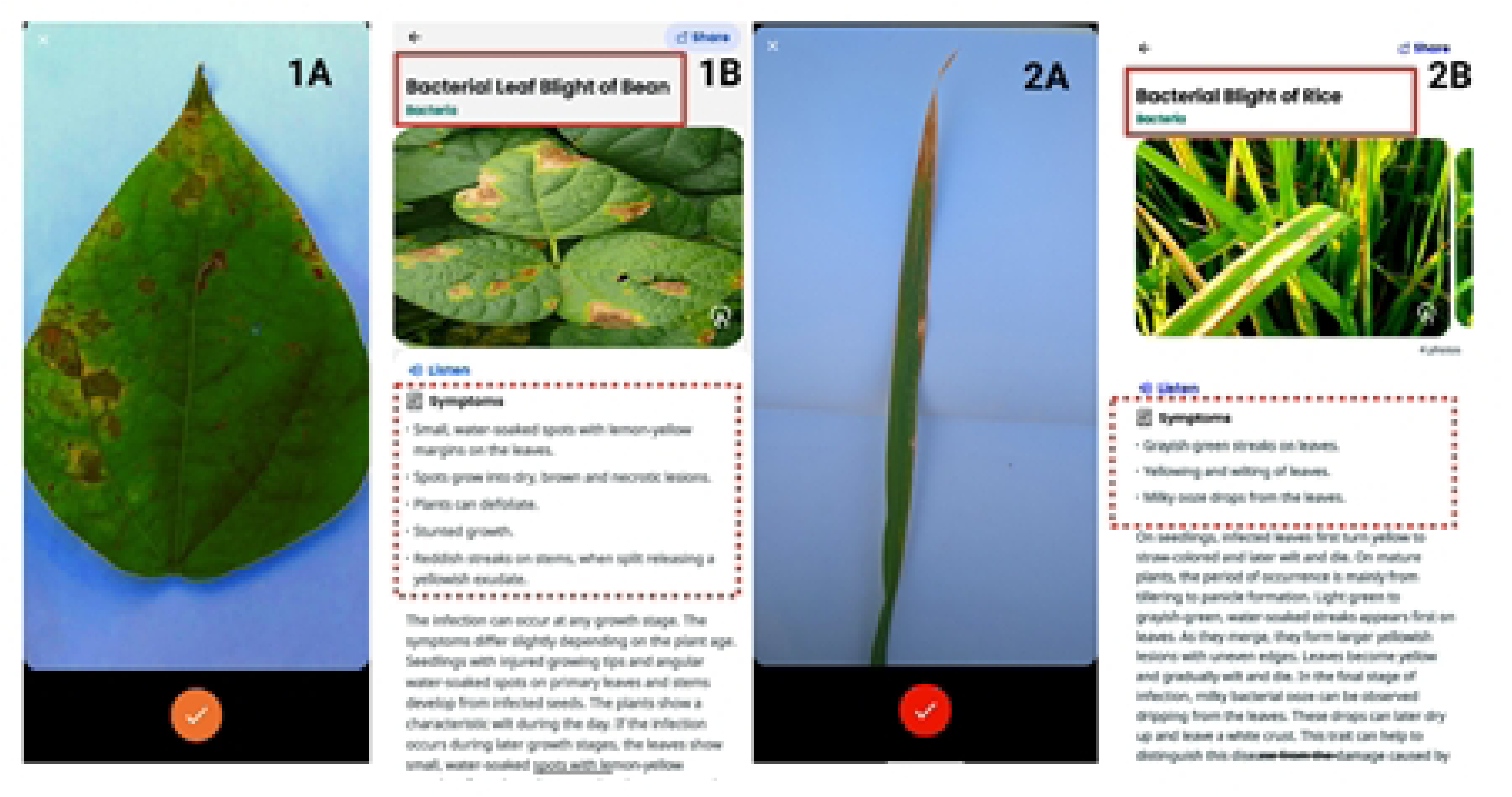
Disease identification steps using Plantix analysis. Illustration of the identification steps of CBB and BLB through the Plantix mobile application. For CBB, image 1A shows the captured leaf sample, while 1B presents the analysis results, including the disease name and associated symptoms. Similarly, for BLB, image 2A displays the captured rice leaf, and 2B shows the corresponding analysis results. In each analysis result image (1B and 2B), a solid red rectangle at the top highlights the identified disease name, while a dotted red rectangle in the middle highlights the associated disease symptoms. Source: Author

#### b) Quantification

After the confirmation of the infection of either CBB or BLB, disease incidences were determined by dividing the number of infected plants by the total number of sampled plants per quadrat, expressed as a percentage, as shown in equation 1.

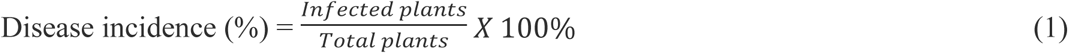

To estimate disease severity, a digital image-based technique was employed. From each quadrat, 25 diseased leaves were sampled for common beans (5 leaves from each of 5 randomly selected infected plants), while 50 diseased leaves were sampled for paddy (5 leaves from each of 10 randomly selected infected hills). Images of both the upper and lower leaf surfaces were captured, giving a total of 125 and 250 diseased leaves per field for common beans and paddy, respectively. Before estimating severity using the captured images, the images were first processed by removing the non-uniform background to create a high-contrast white background. Following the protocol described by [24], the processed images were analyzed in ImageJ, which uses segmentation to annotate and differentiate diseased tissue from healthy tissue (see Fig 4). After segmentation and annotation, disease severity was calculated as the ratio of diseased area pixels to the total leaf area pixels, as shown in equation 2.

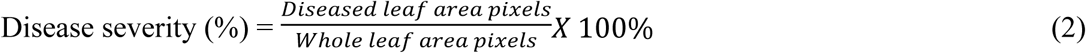

Each leaf’s disease severity (%) was averaged to determine the severity per plant within each quadrat. These plant-level averages were then used to calculate the mean disease severity per quadrat. Finally, the quadrat-level means were subsequently averaged to estimate the overall disease severity for each field.

**Fig 4.**
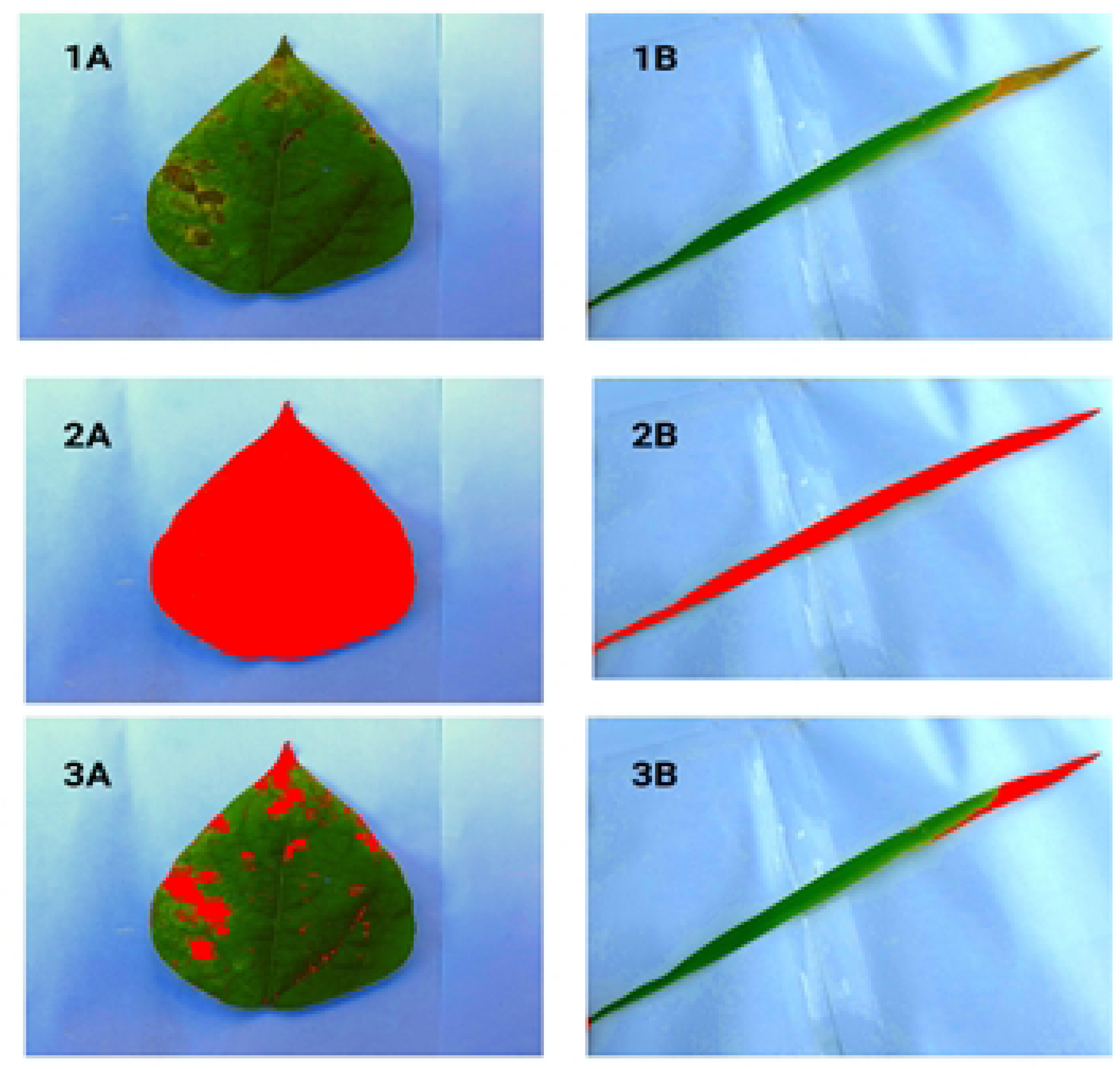
Segmentation steps during ImageJ analysis. Step 1 (1A and 1B): Acquisition of high-resolution images of common bean and paddy leaves infected with CBB and BLB, respectively. Step 2 (2A and 2B): Segmentation of the whole leaf from the background to measure total leaf area. Step 3 (3A and 3B): Segmentation of infected regions from the whole leaf to quantify the diseased area. Source: Author

To estimate disease prevalence, we followed the approach of [6]. Specifically, prevalence was calculated as the proportion of fields affected by CBB or BLB relative to the total number of fields surveyed in each village, as shown in equation 3. The district-level prevalence was then obtained by averaging the prevalence values of all villages within the respective district.

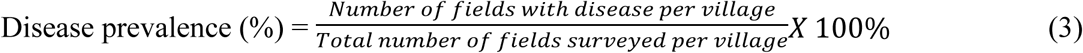

## Data Analysis

Descriptive statistics (i.e., mean percentages and standard deviations) were computed to summarize the distribution of disease prevalence, incidence, and severity across the districts and villages nested within. To assess the variability in the prevalence, incidence, and severity of CBB and BLB across the study districts, a two-level nested Analysis of Variance (Nested ANOVA) was conducted using RStudio (version 4.4.2, R Core Team), with the significance level set at α= 0.05. The analysis considered the districts as the higher-level fixed factor and the village as a lower-level random factor nested within the district, reflecting the hierarchical structure of the data. Before ANOVA, the normality of the disease incidence and severity data was assessed using the Shapiro-Wilk test at a 5% significance level. Variables that did not meet the normality assumption were arcsine-transformed to stabilize variance and improve normality. Since the nested ANOVA does not account for the individual effect of random factors (i.e., villages), a one-way ANOVA was also conducted, treating village as a fixed effect nested within district, with α= 0.05. Where significant differences were detected, post-hoc pairwise comparisons of village means were performed using Tukey’s Honest Significant Difference (HSD) test to identify which villages differed significantly.

To quantify the strength of each factor, effect sizes were calculated using eta-squared (η²). For models with a nested or random structure, partial eta-squared (η²□) was used, as it represents the proportion of variance explained by a factor after accounting for other factors and random effects, providing a measure of the factor’s unique contribution to variation in CBB and BLB incidence and severity. For simple one-way ANOVA models with independent observations, classical eta-squared (η²) was reported, as partial and classical values are identical in this case. Effect sizes (partial eta-squared) were interpreted using the guidelines proposed by [25], as shown in Table 1.

**Table 1:** Interpretation of Partial Eta-squared (η²p) Effect Sizes.

| Value | Effect size |
| --- | --- |
| $X < 0.01$ | Negligible effect |
| $0.01 > X \geq 0.06$ | Small Effect |
| $0.06 > X \geq 0.14$ | Moderate Effect |
| $X > 0.14$ | Large Effect |

X= Classical/ Partial Eta squared value.

## Results

### Incidence, severity, and prevalence of CBB and BLB at the district level

A total of 24 common bean and paddy fields were surveyed for CBB and BLB incidence, severity, and prevalence assessment across Kilosa and Mbarali. CBB was 100% prevalent with varied incidence and severity across both districts. A significantly higher mean incidence (p < 0.005) was recorded in Kilosa (56.80 ± 21.57%) compared to Mbarali (34.31 ± 17.90%) (Table 2; Figure 4A). Similarly, a significantly higher mean severity (p = 0.026) was recorded in Kilosa (15.35 ± 9.74%) than in Mbarali (10.17 ± 4.16%) (Table 2; Fig 5). The results of partial eta squared (η²p), as interpreted by [25], further indicated that large proportions of the spatial variability in incidence (29%) and severity (19%) at the district level were explained by variations among the districts (Table 2).

**Fig 5.**
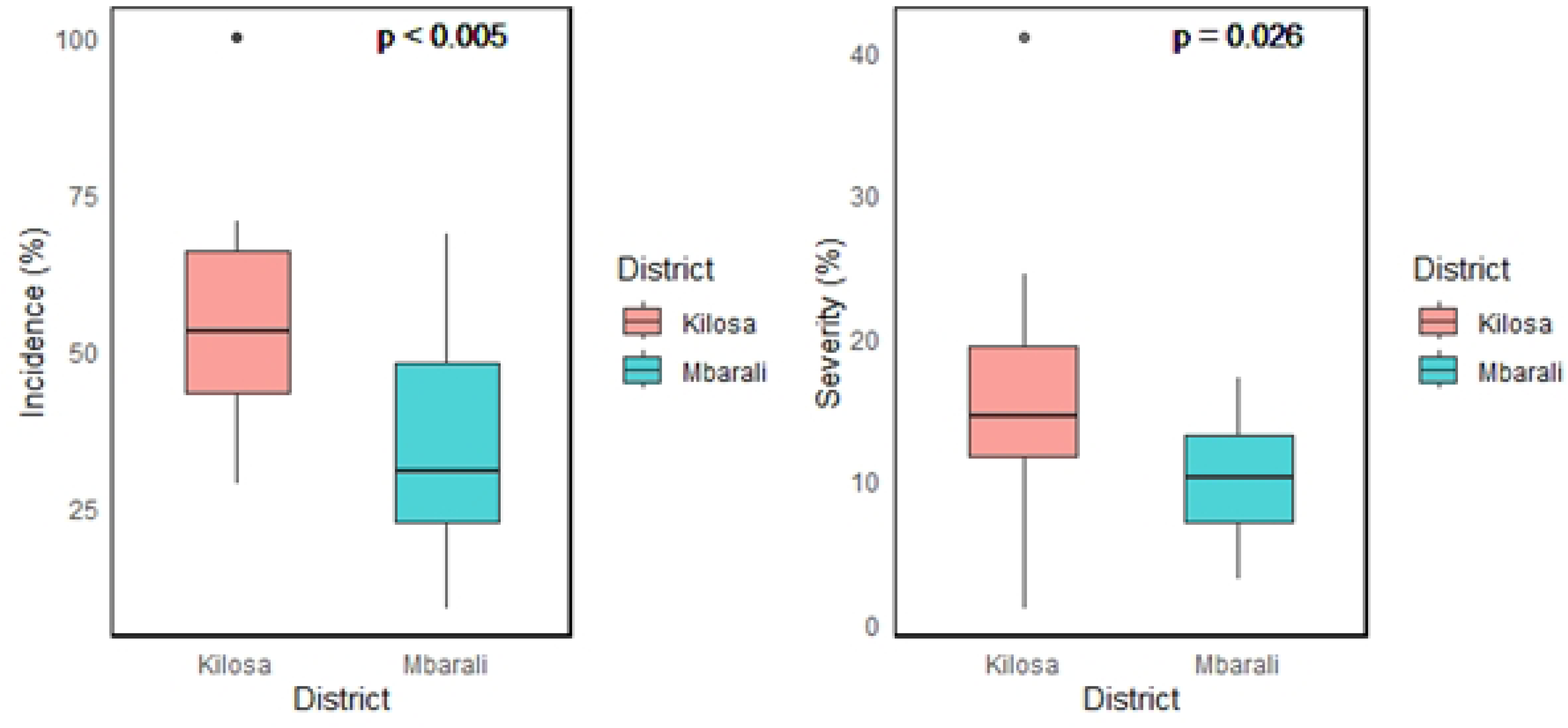
Distribution and comparison of CBB incidence (left) and severity (right) between two study districts. For each district, the thick horizontal line represents the median of the distribution, the box includes 50% of the data, and the whiskers reach the highest and lowest value within 95% of the distribution. Black circles represent outliers outside this range.

**Table 2:**
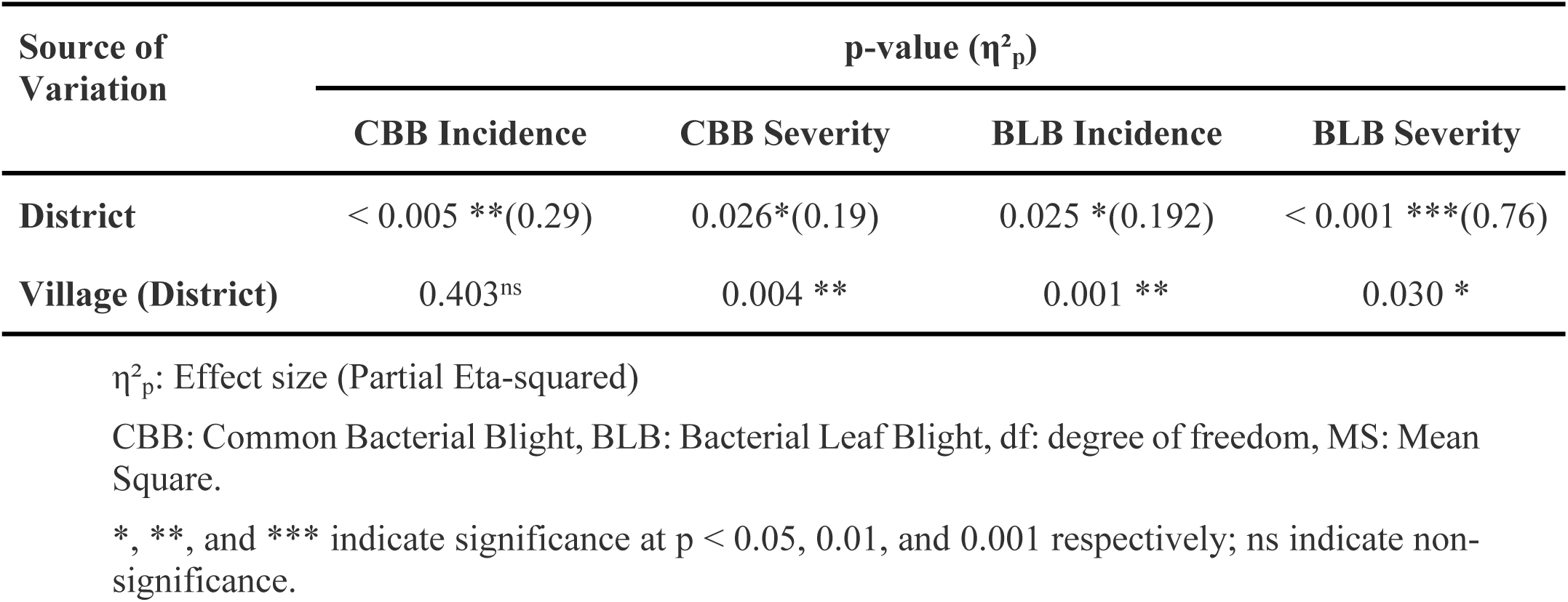
Summary of two-level Nested Analysis of Variance (ANOVA) for the fixed effect (Districts) and random effect (Villages) on the incidence and severity of CBB and BLB, with p-values and Partial Eta-squared (η²□) effect sizes.

BLB was found in all surveyed areas, with a recorded prevalence being 100%. Despite this, incidence and severity differed across the districts. In Kilosa, the overall incidence was 61.46%, which was significantly higher (p = 0.025) than in Mbarali, where the recorded overall incidence was 46.84% (Table 2 and Fig 6). Regarding BLB severity, Kilosa recorded 61.13%, which was significantly higher (p < 0.001) compared to Mbarali (19.3%) (Table 2 and Figure 4B). The effect size analysis (partial eta squared) indicated that variations in incidence and severity were largely explained by differences between the districts, accounting for 19.2% of the variance in incidence and 76% of the variance in severity (Table 2)

**Fig 6.**
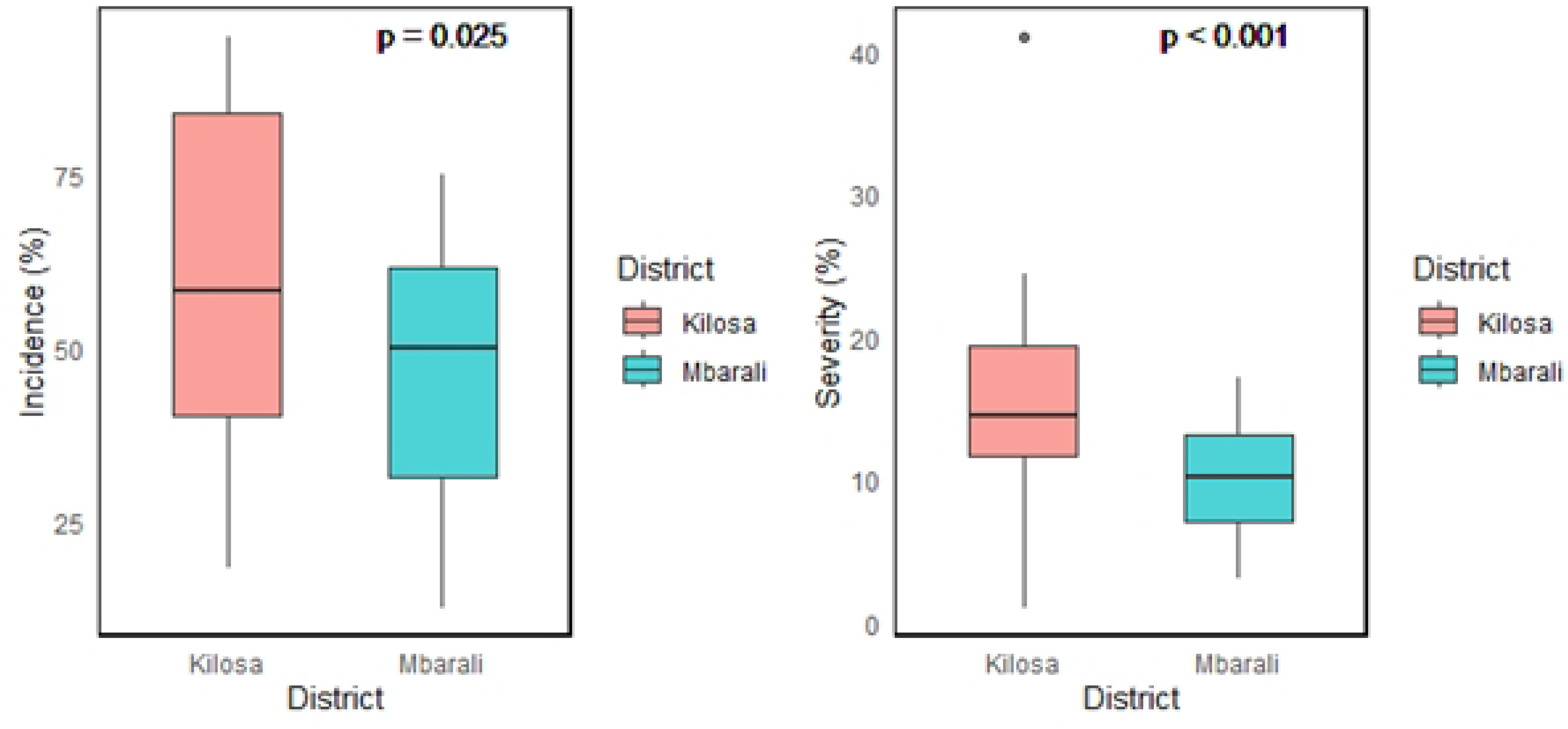
Distribution and comparison of BLB incidence (left) and severity (right) between two study districts. For each district, the thick horizontal line represents the median of the distribution, the box includes 50% of the data, and the whiskers extend to the highest and lowest values within 95% of the distribution. Black circles represent outliers outside this range.

### Incidence, severity, and prevalence of CBB and BLB at the village level

The results of the spatial variability in CBB and BLB incidence and severity at the village level, nested within Mbarali and Kilosa, are summarized in Table 3. For CBB, villages differed significantly only in severity in Kilosa (p = 0.018) (Table 3 and Fig 7), with the recorded severity ranging from 7.3% to 23.43%. A large proportion (49%) of the observed variation in CBB severity was accounted for by village-level differences. No significant variation among villages was detected for CBB incidence in Kilosa, nor for either incidence or severity in Mbarali.

**Fig 7.**
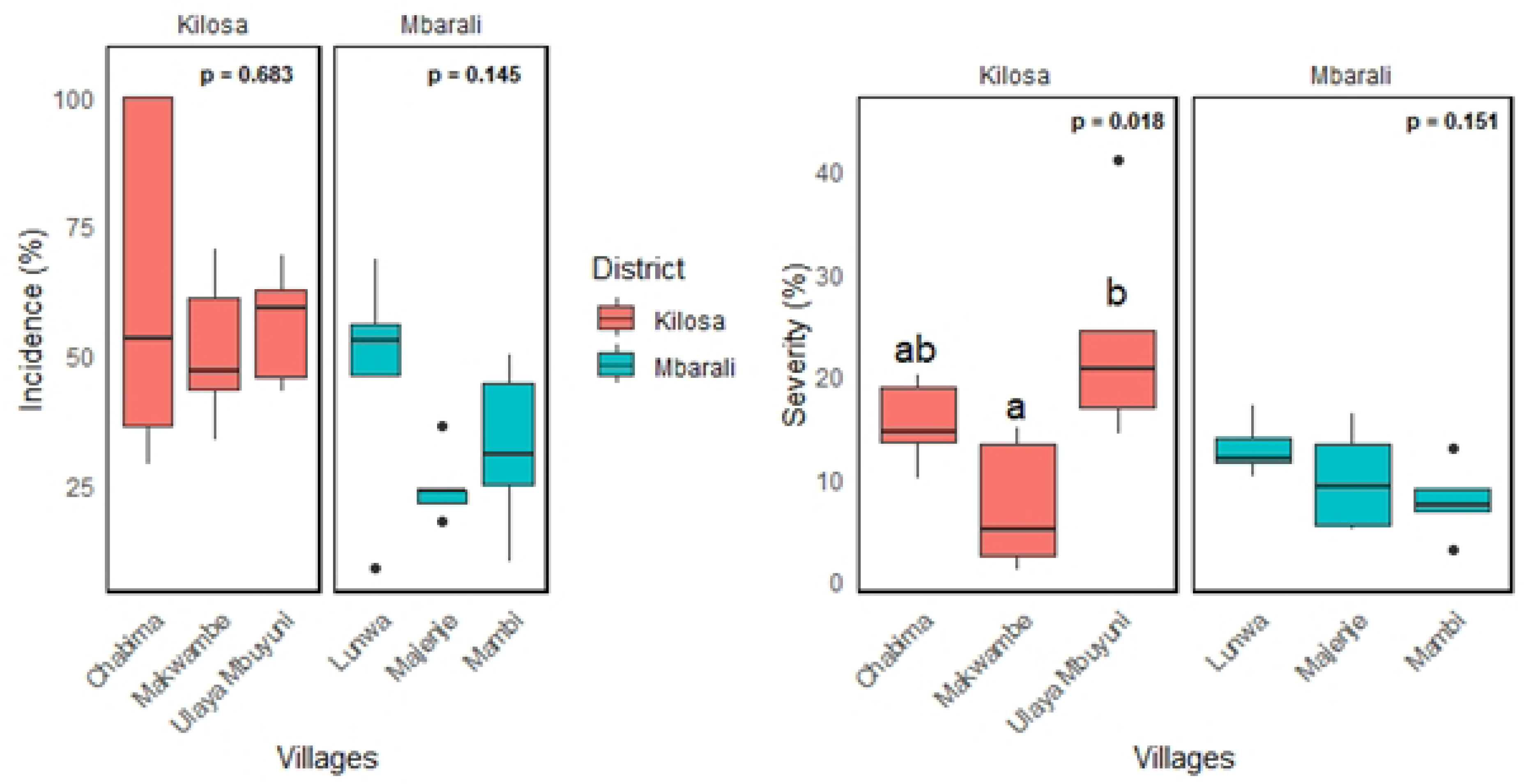
Distribution and comparison of CBB incidence (left) and severity (right) at the village level nested within Kilosa and Mbarali. For each village, the thick horizontal line represents the median of the distribution, the box includes 50% of the data, and the whiskers extend to the highest and lowest values within 95% of the distribution. Black circles represent outliers outside this range. The village’s boxplots bearing different letter(s) differ significantly at the 5% level (See Table 3 for more statistics).

**Table 3:** Effect of random factor (villages) on the incidence and severity of CBB with p-values and Partial Classical eta-squared (η²) effect sizes.

| Villages (Districts) | Incidence (%) | Severity (%) |
| --- | --- | --- |
| **Villages in Kilosa effect on CBB** |  |  |
| Makwambe (Kilosa) | 51.036 | 7.299<sup>a</sup> |
| Chabima (Kilosa) | 63.575 | 15.32174<sup>ab</sup> |
| Ulaya-Mbuyuni (Kilosa) | 55.773 | 23.427<sup>b</sup> |
| Mean | 56.795 | 15.349 |
| L.S.D<sub>(0.05)</sub> | 31.106 | 10.36 |
| p-value | 0.6831<sup>ns</sup> | 0.018* |
| $\eta^2$ | 0.062 | 0.49 |
| **Villages in the Mbarali effect on CBB** |  |  |
| Majenje (Mbarali) | 24.5 | 9.841 |
| Lunwa (Mbarali) | 46.380 | 12.876 |
| Mambi (Mbarali) | 32.043 | 7.796 |
| Mean | 34.30759 | 10.171 |
| L.S.D<sub>(0.05)</sub> | 22.678 | 5.284 |
| p-value | 0.145<sup>ns</sup> | 0.151<sup>ns</sup> |
| $\eta^2$ | 0.275 | 0.27 |
| **Villages in Kilosa effect on BLB** |  |  |
| Mvumi (Kilosa) | 66.582 | 73.393 |
| Ilonga-Msalabani (Kilosa) | 74.676 | 48.083 |
| Ulaya-Mbuyuni (Kilosa) | 43.124 | 61.901 |
| Mean | 61.461 | 61.126 |
| L.S.D<sub>(0.05)</sub> | 30.897 | 22.68 |
| p-value | 0.11<sup>ns</sup> | 0.0899<sup>ns</sup> |
| $\eta^2$ | 0.31 | 0.33 |
| **Village in Mbarali effect on BLB** |  |  |
| Azimio-Mswiswi (Mbarali) | 24.453<sup>a</sup> | 28.349<sup>a</sup> |
| Kongolo-Mkola (Mbarali) | 48.631<sup>b</sup> | 14.572<sup>b</sup> |
| Mambi (Mbarali) | 67.422<sup>c</sup> | 14.962<sup>b</sup> |
| Mean | 46.835 | 19.294 |
| L.S.D<sub>(0.05)</sub> | 10.629 | 11.916 |
| p-value | <0.001*** | 0.0436* |
| $\eta^2$ | 0.867 | 0.41 |

CBB: Common Bacterial Blight, BLB: Bacterial Leaf Blight, LSD<sub>0.05</sub> = Least significant difference at 5%, $\eta^2$ : Effect size (Classical Eta-squared).
\*, \*\*, and \*\*\* indicate significance at $p < 0.05$ , $0.01$ , and $0.001$ respectively; ns indicate non- significance
Means along the same column within a specific category and bearing different letter(s) differ significantly at 5%.

For BLB, both incidence (p < 0.001) and severity (p = 0.044) differed significantly among villages within the Mbarali (Table 3 and Fig 8). Specifically, in Mbarali, the incidence ranged from 24.45% to 67.422%, whereas the severity ranged from 14.572% to 28.349%. The differences among villages explain 86.7% and 41% of the variation in incidence and severity, respectively, which, according to [25], represents a large effect of spatial variability at the village level on the incidence and severity of BLB. In contrast, no significant differences among villages were observed in Kilosa for either parameter.

**Fig 8.**
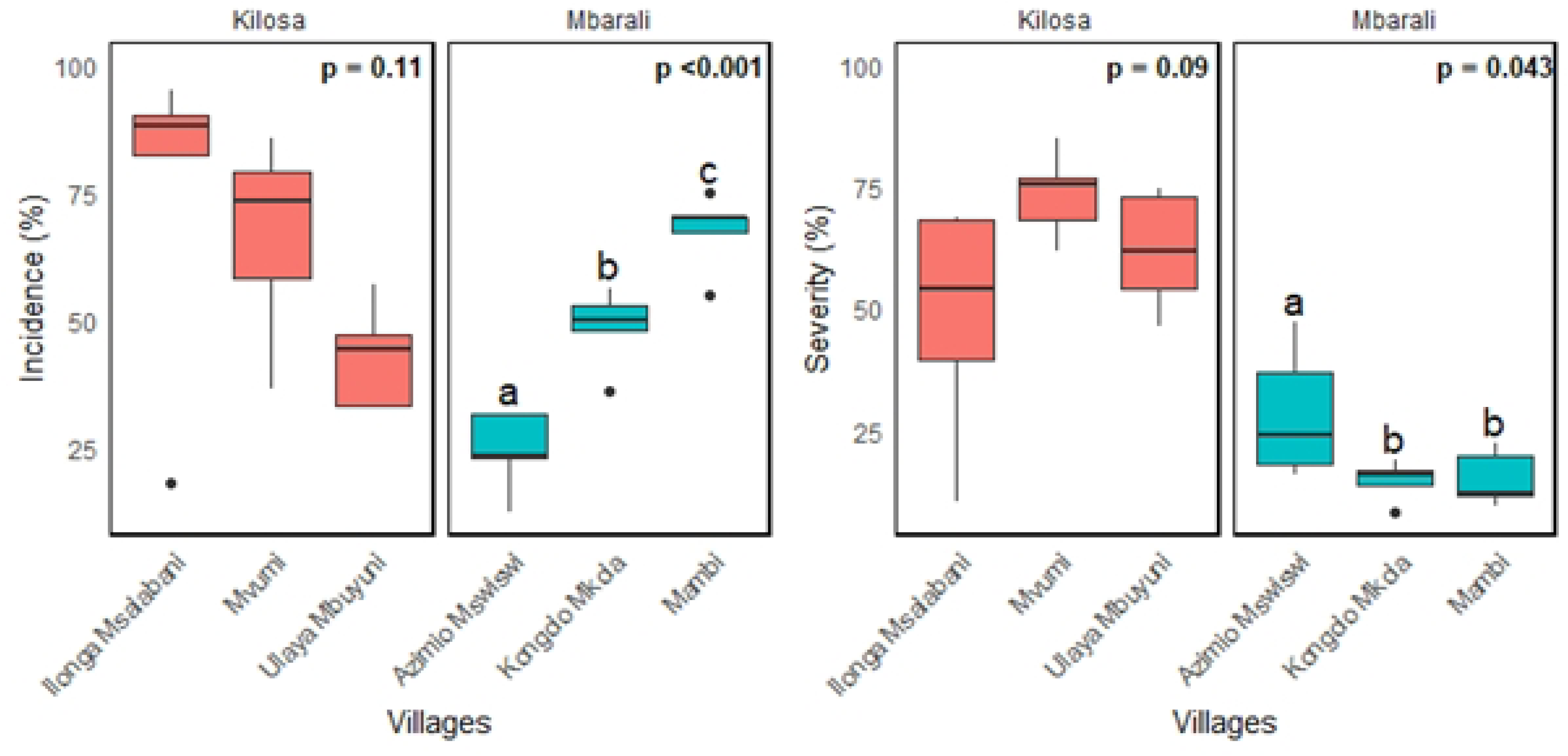
Distribution and comparison of BLB incidence (left) and severity (right) at the village level nested within Kilosa and Mbarali. For each village, the thick horizontal line represents the median of the distribution, the box includes 50% of the data, and the whiskers extend to the highest and lowest values within 95% of the distribution. Black circles represent outliers outside this range. The village’s boxplots bearing different letter(s) differ significantly at the 5% level (See Table 3 for more statistics).

## Discussion

The study aimed to determine the spatial distribution of (CBB) in common beans and (BLB) in paddy across Mbarali and Kilosa districts by assessing prevalence, incidence, and severity using DIA software, which are Plantix and ImageJ. It also aimed to evaluate the spatial variability of these diseases across districts and villages using quantitative measurements.

For both BLB and CBB, higher incidence and severity were recorded in Kilosa than in Mbarali, with clear spatial variation observed at both the district and village levels. Starting at the district level, significantly higher incidence and severity were recorded in Kilosa compared to Mbarali for both CBB and BLB, which indicates a strong spatial heterogeneity of the disease. The significant differences in disease incidence and severity between Kilosa and Mbarali suggest that spatial factors influence the variability of disease intensity across the study area. The effect size analysis, measured using partial eta-squared (η²_p_), further confirmed this, showing that district-level differences accounted for a large proportion of the observed variation in incidence and severity. These differences can be attributed to both environmental conditions and agronomic practices (biophysical factors), which play a critical role in shaping disease epidemics [6,26,27]. First, environmental conditions play a crucial role, as CBB thrives under warm temperatures of 25 to 30 °C, high rainfall, and relative humidity above 80%, which favor bacterial germination, multiplication, and infection [6,28]. Similar to CBB, BLB thrives under temperatures ranging from 25 to 30 °C, high humidity (>70%), and rainfall above 200 mm [29–31]. Since Kilosa is relatively warmer and wetter than Mbarali (see study area section, Figure 2), it provides a more favorable environment for the development and spread of *Xap* and *Xoo*, the causal agents of CBB and BLB, respectively. This could explain the significantly higher incidence and severity of CBB and BLB recorded in Kilosa, whereas the cooler and drier conditions in Mbarali are less conducive to disease progression, thereby limiting its intensity despite high prevalence. Similar to the present findings, other studies have also demonstrated that locations characterized by warm temperatures, high humidity, and substantial rainfall tend to exhibit a higher intensity of CBB and BLB [6,32,33]. Second, differences in agronomic practices, such as variety selection, may have contributed to the observed variation. The adoption of improved resistant varieties of common beans and paddy is widely regarded as a viable and most effective method to manage CBB and BLB [34–36]. However, in Tanzania, it is estimated that more than 70% of farmers rely on farm-saved seeds [37,38]. During the field survey, a similar trend was observed in Kilosa and Mbarali, where most farmers relied on saved landrace seeds for cultivation. Although the resistance profiles of these local varieties are not yet documented, the results suggest that the usage of local landraces may partly contribute to the observed disease intensity in both districts, with the varieties of common beans and paddy grown in Kilosa possessing lower levels of resistance compared to those cultivated in Mbarali, thereby contributing to the higher incidence and severity as recorded in Kilosa. This explanation is further supported by studies such as [6,30], who similarly reported a strong association between the source and type of varieties and the occurrence of CBB and BLB epidemics, respectively. Moreover, since the rapid spread of the severe bacterial blight outbreak, specifically BLB, was first reported in Morogoro [7], where Kilosa district is located, this further confirms that Kilosa’s biophysical factors provide a more conducive environment for the development and spread of the disease.

At the village level, based on our hypothesis, we expected no significant variation in disease incidence and severity among villages within the same district, owing to their proximity and the assumption that they would share similar environmental conditions and agronomic practices. However, our results revealed significant spatial variation of CBB and BLB intensity that extends up to the village level. In Kilosa, the highest CBB severity was recorded in Ulaya-Mbuyuni. In Mbarali, the highest BLB incidence and severity occurred in Mambi and Azimio-Mswiswi, respectively. Furthermore, the effect size analysis, calculated using eta-squared (η²), showed that village-level differences explain a large proportion of variability observed in incidence and severity, further underscoring the impact of differences in local factors among villages on disease intensity. This finding suggests that even within a shared macroenvironment among villages, localized factors can have a significant impact on plant disease epidemics. Differences in agronomic practices across individual fields, such as irrigation, crop spacing, rotation, and intercropping, can create distinct microclimates that influence pathogen development and spread differently across localities, thus causing the observed variability [39–42]. This explanation is further supported by [6], who reported that agronomic practices, including cropping systems and crop density (spacing), are strongly associated with the intensity of CBB. Similarly, [43] found that improper irrigation practices, such as flooding, favour the spread and intensity of BLB. Moreover, because farmers often rely on saved seeds, which are primarily exchanged or gifted through local social networks [44], their distribution tends to be highly localized. As a result, the varieties planted across villages may differ, with neighboring villages potentially cultivating different types. Since different varieties may differ in their resistance to disease, this could partly explain the observed variation in disease intensity across villages.

Despite the spatial variability in CBB and BLB incidence and severity across the study localities, both diseases remained consistently highly prevalent. The consistently high prevalence implies not only that CBB and BLB are currently infectious but also that they are widely distributed across Mbarali and Kilosa districts and the villages nested within. This suggests that, regardless of the variations in biophysical factors at district and village levels, spatial conditions across the study area continued to strongly favor the epidemics of CBB and BLB, as these variations still fall within the range of conditions that favor the growth and development of *Xoo* and *Xap* in suitable hosts, even while modulating disease incidence and severity. These findings are in line with earlier surveys, which also reported that CBB and BLB are highly prevalent and widely distributed across all major common bean and paddy growing areas of Tanzania, including Mbarali and Kilosa [7,8,45,46]. Interestingly, some previous survey studies (e.g.,[47] disagree with the present study’s findings, specifically reporting the absence of BLB in the surveyed villages of the Kilosa district. This contradictory observation indicates an increasing trend in the distribution of BLB and remains a significant challenge for paddy production in the district.

### Strengths and Limitations of the Study

The strength of this study lies in the integration of DIA tools, namely Plantix and ImageJ, to quantify the intensity of CBB and BLB in Kilosa and Mbarali. This approach enabled rapid and accurate identification and measurement of the diseases, which is crucial as they provide a robust basis for planning and decision-making in their management.

However, the study had a few limitations. Firstly, due to logistical constraints, the number of farmers’ fields surveyed in this study was relatively small compared to similar plant disease quantification studies. This may have led to either overestimation or underestimation of disease intensity in the study area, and therefore, the findings should be generalized with caution. Secondly, the study did not identify or quantify the underlying spatial factors responsible for the variability in disease intensity between the study areas. Factors such as differences in soil characteristics, macro- and micro-climatic conditions, crop varieties grown, and local agronomic practices could have influenced the observed patterns but were not explicitly examined. Finally, while disease identification was supported using the Plantix digital diagnostic tool, laboratory-based confirmation of the causal pathogens was not performed, which could have provided additional accuracy and validation of the field diagnoses. Moreover, pathogen-related factors, including the presence of different strains, were not considered, which may have contributed to the observed variability in disease incidence and severity across districts and villages.

## Conclusion

The study revealed that CBB and BLB are highly prevalent across both Kilosa and Mbarali districts and their respective villages; however, their incidence and severity are higher in Kilosa (lowland agroecology) than in Mbarali (highland agroecology), which are strongly associated with the spatial variations between the localities. Generally, this implies that, alongside spatial variations in infection levels, CBB and BLB are highly infectious and widespread in both areas, posing major constraints to the productivity of common beans and paddy. Therefore, improving productivity requires the development of effective, environmentally friendly, and economically viable management strategies that integrate knowledge of disease distribution with the spatial biophysical variability influencing epidemic development at both district and village levels. For farmers, this means that management practices should be tailored to the specific conditions of their fields rather than relying on generic control measures. For researchers, extension services, and relevant authorities, the findings underscore the need for geographically targeted interventions that account for local variability to ensure efficient resource allocation. Accordingly, these efforts should prioritize areas producing common beans and paddy of higher CBB and BLB infection risk, particularly lowland agroecological zones, including Kilosa, while maintaining preventive and surveillance measures in highland agroecological zones, such as Mbarali. Such spatially informed approaches will enhance the efficiency of disease control, reduce yield losses, and strengthen the resilience of common bean and paddy production systems in the face of climate change.

## Acknowledgments

The authors gratefully acknowledge the District Agricultural, Livestock, and Fishery Officers (DALFOs) of Mbarali and Kilosa for providing essential logistical support and guidance during the farm field survey. Their assistance in coordinating with local leaders and facilitating access to the surveyed farms was vital to the success of this study.

